# Convergent evolution of *Streptomyces* protease inhibitors involving a tRNA-mediated condensation-minus NRPS

**DOI:** 10.1101/2020.10.26.356543

**Authors:** César Aguilar, Karina Verdel-Aranda, Hilda E. Ramos-Aboites, Marco Antonio Morales, Cuauhtémoc Licona-Cassani, Francisco Barona-Gómez

## Abstract

Small peptide aldehydes (SPAs) with protease inhibitory activity are natural products typically synthesized by nonribosomal peptide synthetases (NRPS). SPAs are widely used in biotechnology, as therapeutic agents, they are physiologically relevant and regulate development of the natural hosts. During genome evolutionary analysis of *Streptomyces lividans* 66 we identified an NRPS-like biosynthetic gene cluster (BGC) that lacked a condensation (C) domain but included a tRNA-Utilizing Enzyme (tRUE) belonging to the leucyl/phenylalanyl (L/F) transferase family. This system was predicted to direct the synthesis of a novel SPA with protease inhibitory activity, called livipeptin. Following genome mining and phylogenomic analyses we confirmed the presence of tRUEs within diverse *Streptomyces* genomes, including fusions with a C-minus NRPS-like protein. We further demonstrate functional cooperation between these enzymes and provide the biosynthetic rules for the synthesis of livipeptin, expanding the known universe of acetyl-leu/phe-arginal SPAs. The L/F-transferase C-minus NRPS productive interaction was shown to be tRNA-dependent after semisynthetic assays in the presence of RNAse, which contrasts with leupeptin, an acetyl-leu-arginal SPA that we show to be produced by *Streptomyces roseous* ATCC 31245 via a tRUE-minus BGC with multiple complete NRPSs. Thus, livipeptin and leupeptin are the result of convergent evolution, which has driven the appearance of unprecedented biosynthetic logics directing the synthesis of protease inhibitors thought to be at the core of *Streptomyces* colony biology. Our results pave the way for understanding this *Streptomyces* trait, as well as for the discovery of novel natural products following evolutionary genome mining approaches.

**Abstract importance:** Convergent evolution in microbiology is believed to be highly recurrent yet examples that have been comprehensively characterized are scarce. Proteases inhibition by small peptide aldehydes is at the core of many microbiological processes, both within the cell and during colony development, and in microbial ecology. Here we report the biosynthetic foundations of leupeptin, the main Streptomyces protease inhibitor, and of livipeptin, a protease inhibitor produced by Streptomyces lividans. Although these peptides belong to the same chemical class, here we show that their biosynthetic routes result from convergent evolution, as they involve unrelated biosynthetic mechanisms, including the recruitment of a tRNA-utilizing enzyme that functionally replaces the condensation domain of a nonribosomal peptide synthetase during livipeptin biosynthesis. Thus, these results pave the way for understanding Streptomyces protease inhibitors as a trait and provide unprecedented knowledge for genome mining of natural products and synthetic biology where proteases inhibition is desirable.

## Introduction

Small peptide aldehydes (SPAs) are metabolites with protease inhibitory activity. Their production has been reported in several bacterial species belonging to the phyla Actinobacteria, Cyanobacteria and Firmicutes, as well as in fungal species belonging to the families *Aspergillaceae* and *Apiosporaceae* (Sabotič and Kos 2012). SPAs molecular weight ranges between 300 and 900 Da, and they are characterized by unique chemical features, including (i) N-terminal groups capped with acyl groups or with ureido-amino acid groups, giving rise to acylated or aminoalkyl ends with a terminal carboxylic group; and (ii) a terminal aldehyde group derived from the carboxyl terminal modification of the peptide chain by a reductive process (Bullock et al. 1996). Biosynthetically, SPAs have been shown to be produced by nonribosomal peptide synthetases (NRPS) with a reductase (R) domain, which mediates the release of the nascent peptide, and at the same time, the formation of the aldehyde group. These reactions lead to the chemical warhead by which SPAs covalently interact with serine or cysteine proteases active site residues, giving place to hemiacetals or hemithioacetals, and irreversibly inhibiting proteases enzymatic activity (Brayer et al. 1979; Wlodawer et al. 2001).

Based on their functional groups, SPAs can be divided into two sub-classes: (i) those with a terminal group protected by an acyl group, including flavopeptin, tyrostatin, tyropeptin, nerfilin, strepin, leupeptin, thiolstatin, acetyl-leu-arginal and bacithrocins A-D (**Figure 1**); and (ii) those in which the N-terminal group binds to an ureido moiety that is attached to an amino acid by means of an amide linkage, such as chymostatin, MAPI, GE20372, antipain and elastatinal (Umezawa et al. 1970; Suda et al. 1972; Tatsuta et al. 1973; Watanabe et al. 1979; Stefanelli et al. 1995; Murao and Watanabe 2014). The chemical configuration of SPAs allows alteration of the peptide chain, wherein the ureido group acts as an adapter that changes the order of the peptide, from N-terminal to C-terminal into C-terminal to C-terminal, making them distinct from traditional ribosomal peptides. Amongst SPAs, the most widely used is the acetyl-leu/phe-arginal metabolite leupeptin, which was discovered by Umezawa and co-workers more than fifty years ago (Aoyagi et al. 1969; Kondo et al. 1969). Yet, not much is known about the genetic basis and concomitant mechanisms driving the chemical structural diversity and biomolecular specific activities of leupeptin, and overall, of SPAs.

**Figure 1.**
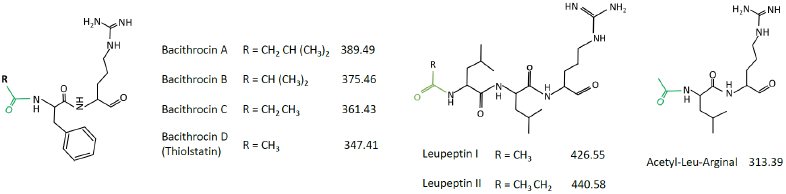
Chemical structures of selected small peptide aldehydes. Structural features of livipeptin (or thiolstatin), and other related SPAs, are shown. The characteristic protecting acetyl group of this family is shown (green).

Previous analysis of the genome of *Streptomyces lividans* 66 and closely related organisms, including *S. coelicolor* M145 and *S. lividans* TK24, led to the detection and further prediction of an unprecedented peptide biosynthetic system, unique to *S. lividans* strain 66 (Cruz-Morales et al. 2013). The predicted biosynthetic system includes (i) an NRPS-like protein, lacking both canonical condensation (C) and thioesterase (TE) domains, but with one adenylation (A) domain, a peptidyl carrier protein (PCP) and a reductase (R) domain (*Sli0883* gene); (ii) a homolog of a leucyl/phenylalanyl (L/F) tRNA transferase (*Sli0884* gene), a tRNA-utilizing enzyme hitherto never implicated in natural products biosynthesis; and (iii) an *N*-acyltransferase (*Sli0885* gene). These enzymes are located within the SLP3 linear plasmid, a mobile genetic element of *S. lividans* absent from the industrial strain TK24 (Cruz-Morales et al. 2013). Our previous proposal implicated these genes in the synthesis of an *N*-acylated leu-arginal dipeptide, which we called livipeptin. Since this prediction suggests common structural features with previously described SPAs, such as leupeptin (Aoyagi et al. 1969; Kondo et al. 1969), thiolstatin (Murao et al. 1985; Kamiyama et al. 1994), bacithrocins (Kamiyama et al. 1994) and acetyl-leu-arginal (Aoyagi et al. 1969; Kondo et al. 1969), it was hypothesized that livipeptin could have protease inhibitory activity.

Here, we characterize *S. lividans* 66 putative livipeptin”s BGC following a combined phylogenomics, genome mining and synthetic biology approach. As predicted, our results show that this unprecedented biosynthetic system directs the synthesis of livipeptin, which shows strong protease inhibitory activity. Livipetin was found to be chemically related to bacithrocins and thiolstatin, reported previously to be produced by *Brevibacillus laterosporus* (Kamiyama et al. 1994); and to leupeptin, whose BGC in *Streptomyces* is also identified herein. Interestingly, leupeptin and livipeptin were shown to be synthesized by unrelated pathways, demonstrating convergent evolution within *Streptomyces* and with other taxonomically unrelated leupeptin-producing proteobacteria (Li et al. 2020). Genetic and semisynthetic chemical experiments in the presence of RNAse, as done previously for other tRUEs (Ortega and van der Donk 2016), confirmed the involvement of an L/F transferase, typically involved in protein tagging during proteolysis (N-end rule) within an essential process carried out by many organisms that regulate the half-life of proteins and morphological differentiation (Leibowitz and Soffer 1969; Varshavsky 2011), during livipeptin biosynthesis.

We anticipate that the discovery of this tRUE, which functionally interacts with a C-minus NRPS, will pave the way for exploiting this unprecedented biosynthetic logic to uncover novel natural products, and will assist in the development of auxiliary synthetic biology tools to inhibit proteolysis (Cruz-Morales et al. 2015a & 2015b). Similar implications are envisaged for leupeptin once its BGC is fully characterized. Moreover, our results are the first step towards investigating the evolution and role of protease inhibitors during *Streptomyces* development, colony biology and ecology, an old hypothesis that remains to be experimentally tested (Chater et al. 2010).

## Results & Discussion

### Genome mining and phylogenomics of actinobacterial L/F transferases

The presence of an L/F transferase homolog in the livipeptin BGC suggested that this enzyme family could have been recruited for the synthesis of a natural product (NP). This possibility encouraged us to mine for L/F transferases in the context of NP biosynthesis, with an emphasis in Actinobacteria, as close homologs of *Sli0884* linked to *Sli0883* could not be found beyond this phylum. Thus, combined EvoMining (Sélem-Mojica et al. 2019) and CORASON (Navarro-Muñoz et al. 2020) phylogenomic analyses of actinobacterial L/F transferases allowed us to identify 137 L/F transferase homologs within 1,246 good quality and well annotated actinobacterial genomes. Remarkably, despite the fact that L/F transferases are known to play a central role as housekeeping enzymes involved in proteolytic metabolism (Mogk et al. 2007), only 11% of the actinobacterial genomes investigated include a homolog of this gene, suggesting that these organisms may have alternative proteolytic tagging strategies, such as the SsrA (tmRNA) tagging system (Braud et al. 2006).

Both the EvoMining and CORASON output revealed two discrete clades, consistent with one function possibly devoted to housekeeping metabolism, and a second function predicted to be involved in NP biosynthesis. The large branch includes 104 enzymes from diverse Actinobacteria species, but without a conserved gene neighborhood (**Supplementary Figure S1**), supporting a single-enzyme housekeeping biochemical function. The second clade is less populated yet shows a larger degree of gene order conservation amongst its 33 entries. These enzymes are mostly from the genus *Streptomyces*, but also from *Nocardiopsis* (1), *Kitasatospora* (3), *Streptacidiphilus* (6) and *Actinopolyspora* (1), with a *Frankia* non-conserved entry at its root (**Figures 2A** and **2B**). This so-called small *Streptomyces* conserved clade consists of L/F transferases probably recruited into NP biosynthesis, as revealed by EvoMining in the first instance, and independently confirmed with the use of antiSMASH (Blin et al. 2019). These analyses also revealed potentially translational fusions between the L/F transferase tRUE and the C-minus NRPS-like genes (shown with an asterisk, **Figure 2B**). As in *S. lividans*, the latter were always found to encode for an adenylation domain predicted to have specificity towards an arginine residue, a peptidyl carrier protein (PCP) and a reductase (R) domain.

**Figure 2.**
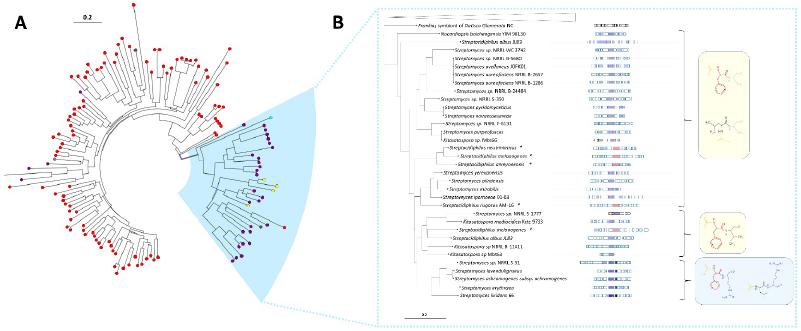
Combined EvoMining and CORASON analysis of L/F transferases in Actinobacteria. **A. EvoMining**. Different metabolic roles or fates for the L/F transferase homologs were predicted: central metabolism (red), transition enzyme (purpule), NP biosynthesis (turquoise), expanded enzymes (grey), and protein fusions with C-minus NRPS-like genes (yellow). The so-called small *Streptomyces* conserved clade is highlighted in blue and analyzed in **B. CORASON**. The large non-conserved clade is used as the root of the small *Streptomyces* conserved clade, which conserves gene neighborhood. L/F transferase (red), C-minus NRPS-like (blue), *N*-acyltransferase (yellow), *O*-methyltransferase (black), butyryl-CoA dehydrogenase (green). An asterisk (*) and pink is used to indicate protein fusions between the C-minus NRPS-like protein and L/F transferases. Structural predictions based on sequence analysis are provided in the right-hand side of the panel.

The domain organization confirmed above resembles that of a canonical NRPS involved in the synthesis of other protease inhibitors (Winn et al. 2016), but with the L/F transferase replacing (and thus possibly playing a similar role to) the expected condensation domain needed for formation of an amide bound. The R domain could be functionally equivalent to that in flavoprotein, an *Streptomyces* SPA of a different class, whose biosynthesis involves an NRPS with a R domain responsible for reductive release of the nascent peptide (Chen et al. 2013). Within the small *Streptomyces* sub-clade, the livipeptin BGC branch includes hits from four other *Streptomyces* species, and one from *Actinopolyspora erythreae* JPMV. Yet, despite these species are from different genera, their BGCs share high sequence similarity and conserved gene order. This includes two other biosynthetically potential elements: an *O*-methyltransferase and a butyryl-CoA dehydrogenase (*Sli0886* and *Sli0887* in *S. lividans*). All together, these observations support our previous prediction related to the synthesis of the putative protease inhibitor livipeptin in *S. lividans* (Cruz-Morales et al. 2013), but also includes leucine in addition to phenylalanine as an alternative biosynthetic precursor (**Figure 3A**).

**Figure 3.**
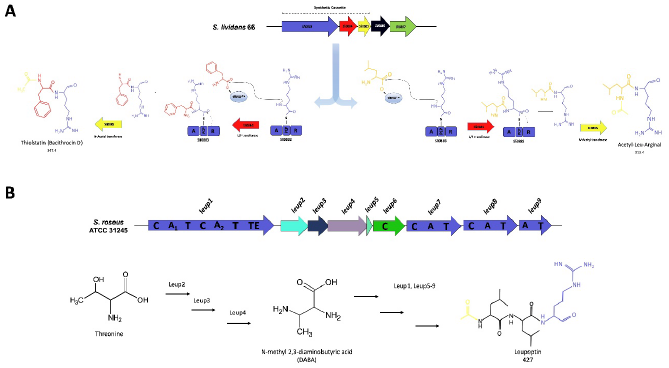
Biosynthetic proposals for livipeptin and leupeptin. **A. Livipeptin biosynthetic pathway.** Dotted lines above the BGC indicate the region included in the synthetic cassette for the heterologous experiments in *E. coli*. Refer to the text for details about livipeptin biosynthetic proposal. **B. Leupeptin biosynthetic pathway**. The postulated BGC directing the synthesis of leupeptin is rich in NRPS genes, in addition to a DABA biosynthetic cassette. Although L-leucine seems the most plausible precursor of leupeptin this has not been confirmed, opening the possibility of DABA serving that role on an uncharacterized pathway.

Owing to the predicted chemical structure of livipeptin we hypothesized it may chemically classify as an acetyl-leu-arginal, which includes other related molecules (Aoyagi et al. 1969; Kondo et al. 1969), the bacithrocins and thiolstatin, previously isolated from *Brevibacillus laterosporus* (Kamiyama et al. 1994), and leupeptin (Aoyagi et al. 1969; Kondo et al. 1969) (**Figure 1**). Therefore, using the L/F transferase as a beacon, we mined the sixty-one *Brevibacillus* spp. genome sequences available at the time of our analyses, without positive results. Likewise, after genome sequencing and mining of the leupeptin-producing strain *S. roseus* ATCC 31245 (Pridham et al. 1965) we were unable to find evidence of a homologous livipeptin BGC. Instead, we identified a BGC rich in NRPSs with A domains specific to diverse aminoacids, such as L-thronine and L-serine (Leup1), L-isoleucine (Leup7) and L-ornithine (Leup9). Additionally, homologs of argininosuccinate lyase, cysteine synthase, and threonine kinase, previously implicated in the biosynthesis of diaminobutyric acid (DABA) were also found (**Figure 3B** and **Supplementary Figure S2**). Although we hypothesized this BGC to be implicated in the synthesis of leupeptin it is unclear how exactly its biosynthesis proceeds.

These observations suggest that despite their chemical similarities, leupeptin and livipeptin BGCs are evolutionarily unrelated, as they follow different biosynthetic logics for their synthesis. While for leupeptin further analysis is need for postulating a complete NRPS biosynthetic pathway, including its biosynthetic precursors (**Figure 3B**), for livipeptin we propose a biosynthetic mechanism involving a small assembly line that catalyzes amide bond formation between arginine (A domain encoded by the NRPS-like gene) and phenylalanine or leucine residues provided by their cognate aminoacyl-tRNAs as substrates (tRUE of the L/F transferase family). A reductive release mechanism leading to the aldehyde moiety of livipeptin would also be part of this pathway. The resulting dipeptide aldehyde may undergo acetylation catalyzed by the function of an *N*-acetyltransferase (*Sli0885*). These three enzymes may represent the minimal biosynthetic core necessary to synthesize a metabolite with protease inhibitory activity. However, additional methylation (*Sli0886*) and further dehydrogenation (*Sli0887*) could also be in place **(Figure 3A)**. In the absence of transcriptional regulatory genes from both BGC, how these enzymes potentially interact to provide a regulatory mechanism controlling protease inhibitory activity, as previously suggested in *Streptomyces* species (Kim and Lee 1995; Kim and Lee 1996), remains unknown.

### S. *lividans* livipeptin and *S. roseus* leupeptin BGCs direct the synthesis of acetyl-leu/phe-arginal metabolites

To identify the livipeptin metabolite(s) potentially produced by *Sli0883-5*, proposed to have protease inhibitory activity, a *S. lividans* 66 mutant was constructed (*S. lividans* ΔLvp, *Sli0883-5* minus; **Table 1**). *S. lividans* strains 66 (wt) and ΔLvp (mutant) grown in modified R5 medium were used for the identification of livipeptin after comparative HPLC analysis of the aqueous extracts of the resulting cultures. These conditions were suboptimal for comparative chromatographic analysis, as they resulted in far too many metabolites, but only one differential peak eluting at a retention time of 4.5 min with a distinctive absorption at 280 nm. The compound associated with this peak could be isolated after extensive HPLC optimization and fractionation (**Figure 4A**), and high-resolution MS analysis of the isolated compound(s) showed that this peak corresponds to a metabolite with an *m/z* of 365.18 (**Supplementary Figure S3**). Further experiments with an MS ion trap targeted toward this mass reproducibly lead to a distinctive fractionation that includes a derived ion with an *m/z* of 347 (**Figure 4B**), equivalent to loss of a water molecule, in addition to ions with *m/z* values of 203 and 185 (**Figure 4C**, see also Methods).

**Table 1.** Chemical and activity features of livipetin from *S. lividans* and leupeptin from *S. roseus*.

| Organism | Genotype | Metabolite | RT<sup>a</sup><br>(min) | MS<sup>a</sup><br>(m/z) | MS<sup>2</sup><br>(m/z) | T. I. A.<sup>c</sup><br>(%) | P. I. A.<sup>c</sup><br>(%) |
| --- | --- | --- | --- | --- | --- | --- | --- |
| *S. lividans* 66 | WT | Livipeptin | 4.5 | 365.18 | 347<br>**203**<br>**185** | 10.5 ± 1.1 | 3.5 ± 0.4 |
| *S. lividans* ΔLvp | WT *Sli0883-5*<sup>-</sup> | Livipeptin | NI | NI | NI | 88 ± 4.1 | 85.4 ± 1.6 |
| *E. coli* pAsk | C41 pAsk<br>(empty vector) |  | NI | NI |  | 87.5 ± 10.9 | 100 ± 6.4 |
| *E. coli* pLvp | *E. coli* C41 pLvp<br>(*Sli0883-84-85*) | Livipeptin | 4.5 | 365.18 | 347<br>**203**<br>**185** | 12.7 ± 0.6 | 11.3 ± 1.8 |
| *S. roseus* ATCC<br>31245 | wild type (WT) | Leupeptin | 28.5 | 427 | 409<br>367<br>365<br>**203**<br>**185** | 9.6 ± 0.7<sup>d</sup> |  |
| *S. roseus* LeuP1 | WT *leupA::kan* | Leupeptin | NI | NI | NI |  |  |
| *E. coli* pLeup | WT pESAC_810<br>(*leup1-9*)<sup>e</sup> | Leupeptin | 28.5 | 427 | 409<br>367<br>365<br>**203**<br>**185** |  |  |

<sup>a</sup> HPLC retention time (RT) and mass spectrometry (MS, MS<sup>2</sup>) data with common fragments between livipeptin and leupeptin shown in bold. NI, not identified (see **Figure 4** and its associated **Supplementary Figures S3** and **S4** for spectral analysis). <sup>b</sup> T.I.A. and P.I.A., trypsin and papain inhibitory activity, respectively, which was recorded as activity curves for 20 minutes, and reported as a percentage of the activity found for leupeptin and papain, respectively (see **Figure 5A**). Data was generated after three independent experiments, leading to standard deviations measured as relative fluorescence units (RFU). <sup>d</sup> Reported activity is from commercial leupeptin of microbial source with ≥ 90% purity (Sigma, No. L2884). <sup>e</sup> pESAC\_810 construct (or pLeup) is a pESAC13-A derivative (Tocchetti et al. 2018) containing the predicted leupeptin BGC, including the *leup1-9* genes.

**Figure 4.**
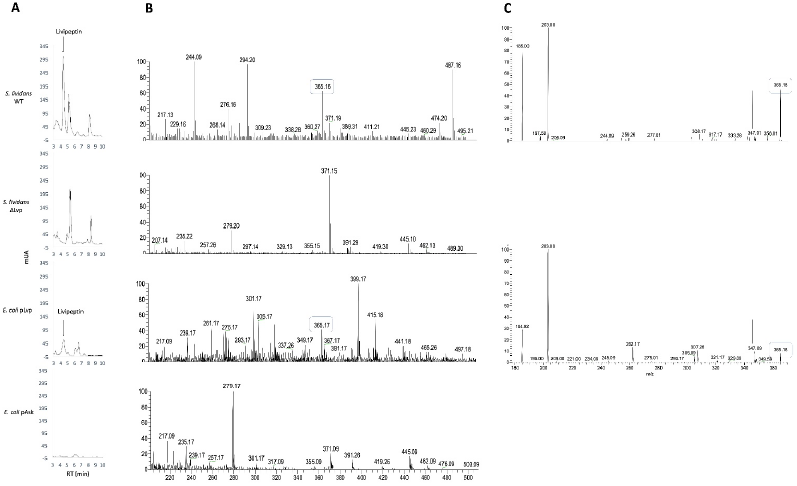
HPLC-MS profiles of homologous and heterologous livipeptin. **A**. HPLC profiles of livipeptin produced by *S. lividans* (WT and mutant, 66 and ΔLvp, respectively) and *E. coli* (overexpressing and empty vector, pLvp and pAsk, respectively). **B**. MS analysis of the purified fractions from **A** eluting at 4.5 mins (280 nm), corresponding to a compound with a mass of 365.18 (*m/z*). Livipeptin”s high resolution MS data is provided as **Supplementary Figure S3. C**. MS^2^ analysis of the peak from **B** produced by *S. lividans* 66 and *E. coli* C41 Lvp sustaining the proposal of livipeptin structure with and without a molecule of water (347 and 365 *m/z*, respectively). A common identity for homologous and heterologous livipeptin is confirmed by the 185 and 203 (*m/z*) fragments. The negative spectra obtained for strains *S. lividans* ΔLvp (mutant) and *E*.*coli* C41 pAsk (empty vector) is provided as **Supplementary Figure S3**.

The abovementioned compound is close to the theoretical molecular formula of thiolstatin (C_17_N_5_O_3_H_21_), but only if incorporation of L-phenylalanine over L-leucine, together with L-arginine, is considered. Indeed, MS^2^ data obtained coincides with previously reported data for other acetyl-leu-arginal metabolites (Aoyagi et al. 1969; Kondo et al. 1969), the bacithrocins and thiolstatin produced by *Brevibacillus laterosporus* (Kamiyama et al. 1994). Moreover, although more distantly, these data confirm the expected chemical connection via leucine residues between livipeptin with leupeptin (185 and 203 fragments, see **Table 1**). Therefore, we concluded that livipeptin is equivalent to thiolstatin and bacithrocin D, despite our inability to identify a homologous locus in the currently available *Brevibacillus* genome sequences. This could be due to the lack of taxonomic resolution and detailed information about the *Brevibacillus laterosporus* strain (Kamiyama et al. 1994), leading to misclassification of the original thiolstatin- and bacithrocin-producing strain(s). After genome mining analyses (data not shown) we favor this explanation rather than the existence of an analogous livipeptin BGC in species belonging to *Brevibacillus*, although a different biosynthetic route yet to be identified cannot be ruled out.

In order to identify the leupeptin BGC, we first confirmed by HPLC-MS the ability of *S. roseus* ATCC 31245 to produce this metabolite. The metabolite found was consistent with the leupeptin standard from a commercial source. We then sequenced and mined the genome of this organism, which consists of 7.8 Mb encoding for 6,765 ORFs and 27 putative BGCs. As mentioned previously, our efforts failed to identify a livipeptin BGC in this streptomycete. Yet, we identified alternative BGCs that could account for the synthesis of leupeptin (Cruz-Morales et al. 2015; Cruz-Morales, et al. 2016). A combined EvoMining, CORASON and antiSMASH analysis gave rise to a hypothesis that a BGC with five NRPSs and a diaminobutyric acid (DABA) biosynthetic cassette (**Supplementary Figure S2**) could direct the synthesis of leupeptin. Full functional characterization of this BGC is beyond the scope of this report, but to confirm the involvement of this locus in leupeptin biosynthesis we constructed an insertional mutant by targeting the *leupA* gene with a suicidal plasmid, leading to a strain termed *S. roseus* LeuP1 (see **Table 1** and **Methods**). This strain was unable to synthesize leupeptin when compared with the wild type parental ATCC 31245 strain. Moreover, heterologous expression of pESAC13-A constructs (Tocchetti et al. 2018) bearing the putative leupeptin BGC in *E. coli* lead to production of leupeptin. The MS^2^ fragmentation pattern of isolated leupeptin coincides with a commercial standard and with some fragments of livipeptin spectral data (185 and 203, see **Table 1** and **Supplementary Figure S4)**.

Our results therefore establish a chemical relationship between livipeptin and leupeptin as acetyl-leu/phe-arginal metabolites, despite involving distinct biosynthetic pathways, and potentially different precursors. While leupeptin is proposed to be synthesized by nine genes (including five NRPSs and a subcluster potentially involved in the synthesis of the non-proteinogenic amino acid DABA), livipeptin is produced by a C-minus NRPS-like protein functionally linked with a tRUE L/F transferase, plus an acyltransferase. On one hand, the identification of distinct biosynthetic pathways to produce very similar NPs highlights the increasing versatility observed among NRPS-derived natural products in terms of their biosynthetic logics and enzyme interactions within multifunctional protein templates. On the other hand, the occurrence of convergent evolution in natural products biosynthesis is a recurring theme (Fischbach 2009), although its significance and rate remains to be determined (Chevrette et al. 2020). It is nevertheless tempting to speculate that the degree of convergent evolution correlates with chemical scaffolds that contribute towards the fitness of the host organisms, as previously suggested for protease inhibitors (Chater et al. 2010; Guo et al. 2017; Li et al. 2020)

### Heterologous livipeptin and leupeptin have protease inhibitory activity

To further characterize livipeptin, and to be able to determine its presumed protease inhibitory activity, we used as positive controls commercial leupeptin and antipain against trypsin and papain, respectively. For leupeptin, three different sources with identical results were used: (i) commercial standard produced after bacterial fermentations, (ii) *S. roseus* ATCC 31245 and (iii) *E. coli* cultures expressing pESAC13-A constructs (Tocchetti et al. 2018), namely pESAC_810, harboring the presumed leupeptin BGC (**Table 1**). For livipeptin, we opted for the use of synthetic *Sli0883-5* genes optimized for *E. coli* and their expression from the pAsk vector (Adams et al. 2014). The resulting expression plasmid, pLvp, was used to transform *E. coli* C41. The resulting strain, C41 Lvp, was cultivated in minimal medium. The conditions used for these experiments allowed us to easily perform a chromatographic comparative analysis of aqueous extracts, without the residual metabolites found in the *S. lividans* native system.

As above, comparison of the HPLC profiles of *E. coli* expressing pLvp and *E. coli* bearing the empty plasmid pAsk showed a differential peak at 4.5 min absorbing at 280 nm **(Figure 4A)**. This metabolite shows up in the *E. coli* cultures with less background than that seen in the *S. lividans* samples (**Figures 4B** and **4C**). Yet, this *E. coli* data also failed to identify the presumed L-leucine livipeptin analog with a mass of 313.39 (*m*/*z*), but did rule out the WT *S. lividans*-specific mass 487 (*m/z*) as related to livipeptin, given that it could not be detected within the *E. coli* pLvp spectra. Whether this mass corresponds to a bigger version of livipeptin was further ruled out after inspection of the MS^2^ spectra of the 487 (*m/z*) metabolite (**Supplementary Figure S3**). Thus, although these data do not confirm that all three *Sli0883-5* genes are needed for synthesis of livipeptin, it unequivocally establishes a link between the *lvp* genes *Sli0883-5* and a putative SPA with protease inhibitory activity.

Previous data suggests that thiolstatin has weak inhibitory activity against serine proteases, such as trypsin, but potent proteolytic inhibitory activity towards cysteine proteases, such as papain. These observations actually explain the use of the prefix “thiol” in its name (Murao et al. 1985). Unfortunately, the absence of thiolstatin or bacithrocins standards hampered our ability to compare the activity of these metabolites with that of livipeptin. Thus, we focused on HPLC-purified products from *S. lividans* 66 and *E. coli* C41 pLvp cultures. The results obtained from these experiments are included in **Table 1** and shown in **Figure 5A**.

**Figure 5.**
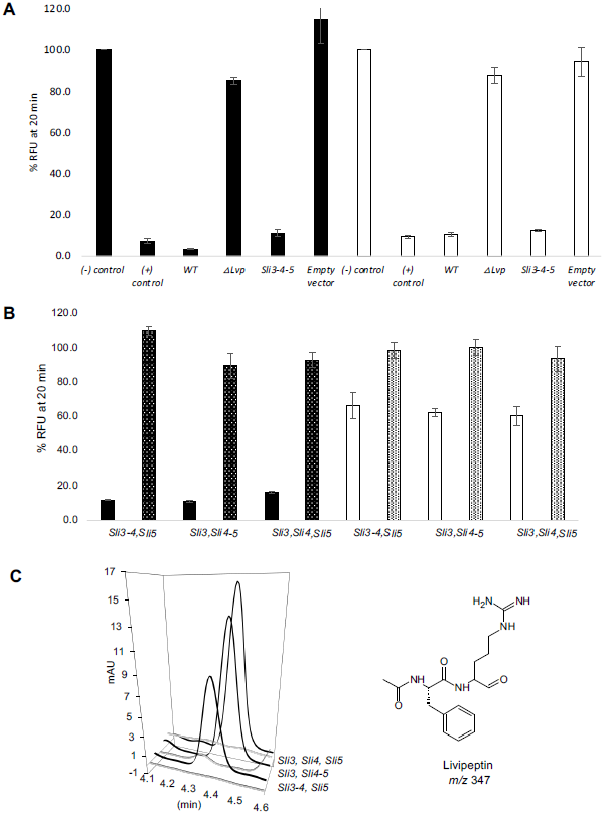
Papain and trypsin inhibitory activity of livipeptin. **A**. Enzyme inhibition assays were performed against papain, with antipain as a control (black bars), and trypsin, with leupeptin as a control (white bars). Results for both native (*S. lividans* WT and ΔLvp mutant) and heterologous (pLvp and empty vector) systems are shown. For the sake of clarity, *Sli0883-5* genes are shown only with their final digit, i.e. 3, 4 and 5. A comma (,) and a hyphen (-) are used to denote expression *in trans* or *in cis*, respectively. Standard deviations shown were calculated from three independent experiments (**Table 1**) **B**. Activity reconstitution using a synthetic biology approach of livipeptin BGC. Metabolites, proteases and controls are as in panel A (data not shown). Only the three three-genes rection mixtures, irrespective of their genetic organization (expression *in trans* or *in cis*), showed protease inhibitory activity. Cell-free extracts with one- or two-genes constructs did not show inhibitory activity (**Table 2**). RNase (dotted bars) eliminates trypsin and papain inhibitory activity in around 87% and 37%, respectively. **C**. Reconstitution of livipeptin (*m/z* 347) biosynthesis used for experiments of panel **B**, showing a tRNA catalytic dependency (first trace of each dataset).

**Table 2.** Reconstitution of livipeptin biosynthesis and its protease inhibitory activity.

| Reaction Mix<br>(*E. coli* c. f. e.)<sup>a</sup> | Genotype<br>(plasmids)<sup>b</sup> | RT<br>(min) | T. I. A.<br>(%) | P. I. A.<br>(%) |
| --- | --- | --- | --- | --- |
| pAsk | Empty vector | NI | 87.5 ± 10.9 | 100 ± 6.4 |
| pLvp | *Sli0883-4-5* | 4.5 | 12.7 ± 0.6 | 11.3 ± 1.8 |
| + RNase |  | NI | 98.7 ± 6.9 | 99.8 ± 8.2 |
| Sli3, pSli4, pSli5 | *Sli0883, Sli0884, Sli0885* | 4.5 | 60.5 ± 5.5 | 16.4 ± 0.4 |
| + RNase |  | NI | 93.9 ± 7.2 | 93.2 ± 4.4 |
| Sli3, Sli4-5 | *Sli0883, Sli0884-5* | 4.5 | 62.5 ± 2.5 | 11 ± 0.5 |
| + RNase |  | NI | 100 ± 4.6 | 89.9 ± 7.3 |
| Sli3-4, Sli5 | *Sli0883-4, Sli0885* | 4.5 | 66.8 ± 7.7 | 12 ± 0.4 |
| + RNase |  | NI | 98.9 ± 4.7 | 100 ± 2.1 |
| Sli3-4 | *Sli0883-4* | NI | 99.2 ± 9.1 | 98.1 ± 7.9 |
| Sli4-5 | *Sli0884-5* | NI | 90.4 ± 9.0 | 100 ± 4.7 |
| Sli3 | *Sli0883* | NI | 100 ± 8.2 | 100 ± 3.1 |
| Sli4 | *Sli0884* | NI | 100 ± 5.9 | 100 ± 4.4 |
| Sli5 | *Sli0885* | NI | 92.5 ± 5.4 | 100 ± 4.6 |

<sup>a</sup> The reaction mixtures were prepared with *E. coli* cell-free extracts (c. f. e.) with or without addition of RNase. <sup>b</sup> Genotype provided is for plasmids harboring the *S. lividans* 66 genes *Sli0883-5*. NI, livipeptin could be not identified. As in **Figure 5**, *Sli0883-5* genes are shown only with their final digit, i.e. 3, 4 and 5. A comma (,) and a hyphen (-) are used to denote expression *in trans* or *in cis*, respectively. Trypsin (T. I. A.) and papain (P. I. A.), HPLC and MS analysis, was determined and expressed as in **Table 1**.

Although strong inhibitory activity equivalent to leupeptin and antipain could be found for the HPLC-purified metabolite(s) from the two experimental sources investigated, the material obtained from the *E. coli* heterologous system showed less activity, especially towards papain. This may contradict early data on thiolstatin activity (Murao et al. 1985; Kamiyama et al. 1994), but given that the overall conditions of these experiments are not equivalent, this is an unfair comparison and we cannot explain this difference. Interestingly, as can be seen in **Figure 4**, the extracts from *S. lividans* 66 have a larger metabolite diversity than that from *E. coli*. Similar to leupeptin (Suzukake et al. 1980), under the rich *Streptomyces* growth conditions, livipetin could be produced and co-purified as a molecular cluster, leading to synergistic protease inhibitory activity. A similar situation has been previously shown for other biomolecular activities of natural products (Gutiérrez-García et al. 2017). Indeed, commercial high quality leupeptin isolated from *S. roseus* cultures includes several related species, which contrasts with the less active synthetic leupeptin (McConnell et al. 1990). Indeed, the HPLC-purified metabolite(s) obtained from the *S. lividans* mutant ΔLvp shows some residual inhibitory activity against both proteases (**Figure 5A**).

### Livipeptin biosynthesis involves a C-minus NRPS tRNA-mediated biosynthetic logic

In addition to codon usage optimization for *E. coli* expression, the pLvp plasmid was designed such that each of the three *lvp* genes (*Sli0883-85*) could be excised and the plasmid circularized with different-yet-compatible cohesive restriction enzyme, easing construction of expression plasmids with all possible combinations (**Supplementary Figure S5)**. In addition to pLvp (pSli3-4-5), as shown in **Table 2**, five different constructs were obtained: three single-gene plasmids (pSli3, pSli4 and pSli5) and two double-gene plasmids (pSli3-4 and pSli4-5). The resulting plasmids were confirmed after DNA sequencing and used in assays consisting of mixtures of different cell-free extracts expressing the plasmids, such that all *cis* and *trans* gene-expression possible combinations could be addressed. This synthetic biology strategy allowed us to evaluate the potential functional impact that co-expression of two different enzymes could have, when compared with independent enzyme expression, providing a sense of the predicted enzyme-enzyme interactions between the C-minus NRPS-like protein, the L/F-transferase and the *N*-acetyl transferase, and its relationship with protease inhibitory activity (**Figure 5B** and **5C**).

These experiments showed that only the three-genes combinations, obtained after independent expression (*in trans*) from independent plasmids or co-expression (*in cis*) from the same plasmid, yielded livipeptin synthesis (**Figure 5C**). Unexpectedly, *in trans* expression of *Sli0883, Sli0884* and *Sli0885* provided the highest yield of livipeptin. The combination *Sli0883,5* and *Sli0884* was not tested as the former plasmid was not obtained, whereas all combinations with only two or one gene did not lead to production of livipeptin (**Table 2**). More importantly, as expected, reconstituted livipeptin showed protease inhibitory activity, similar to that found in the *S. lividans* native system (**Figure 5A**) and the heterologous *E. coli* system (**Figure 5B**) against trypsin, and to a lesser extent, papain. Although the reason of these differences remains unknown, these results establish the *Sli0883-5* genes as the minimal biosynthetic core for the synthesis of livipeptin with protease inhibitory activity.

This experimental layout also provided us with the opportunity to test the hypothesized aminoacyl-tRNA dependency of the L/F transferase tRUE. *In vitro* synthesis of livipeptin with the gene combinations containing the entire cluster was redone. However, for these experiments, the cell-free extracts were pre-incubated with RNase prior to generating the reaction mixture. As shown in **Figure 5C** and **Table 2**, HPLC analysis shows that the addition of RNase suppresses the formation of livipeptin, which can be explained by the degradation of the aminoacyl-tRNA substrate. This approach has been previously adopted during characterization of the tRNA-dependent lantibiotic dehydratase NisB, involved in nistatin biosynthesis (Ortega et al. 2014). As we could not detect protease inhibitory activity in these cell-free extracts, *Sli0883-5* genes are strictly necessary for livipeptin synthesis. These results add to the increasing number of reports that implicate central metabolism tRUEs in natural products biosynthesis (Hong et al. 2004; Garg et al. 2008; Zhang et al. 2011; Belin et al. 2012; Bougioukou et al. 2013; Ortega et al. 2014).

The small size of livipeptin biosynthetic pathway is an outstanding feature of this BGC, which seems only possible due to the catalytic versatility of the tRNA L/F-transferase, predicted to functionally replace the C domain of the NRPS-like protein. Recent studies on tRUEs have focused on protein structure and conservation of the enzyme family (Ichetovkin et al. 1997; Watanabe et al. 2007; Fung et al. 2011; Zhang et al. 2011; Ortega et al. 2014), the catalytic mechanism and residues involved in substrate-assisted catalysis (Watanabe et al. 2007; Fung et al. 2011; Zhang et al. 2011) and in the innovative role of tRNA-dependent enzymes in pacidamycin and lantibiotic biosynthetic pathways (Zhang et al. 2011; Ortega et al. 2014). However, the role of an L/F tRNA transferase in peptide bond formation during biosynthesis of a SPA, in association with an NRPS-like protein, has not been shown until now. The combinatorial potential provided by the interaction between A domains and tRUEs warrants further investigation in many ways. For instance, we envisage exploitation of this discovery for untapping novel natural products diversity through genome mining efforts, as well as for the development of synthetic biology approaches targeting proteolysis in different settings (Hines et al. 2008), such as within the present Covid19 pandemic crisis.

## Materials and Methods

### Phylogenomics and genome mining of natural products

Enzymes from four different microorganisms (*Streptomyces lividans, Rubrobacter xylanophilus, Frankia alni* and *Propionibacterium acnes*), together with the L/F transferase homolog from the livipeptin BGC, were used as query sequences to retrieve homologs from our 1,246 actinobacterial genome database. From this search, 140 homologous sequences were obtained and used to look for known NPs biosynthetic pathways recruitments in the MIBiG database (Kautsar et al. 2019).The BGC for livipeptin and leupeptin were curated and deposited in this robust community standard database under the accession numbers BGC0001168 and BGC0002095 for livipeptin and leupeptin, respectively. The 140 L/F transferase sequences were aligned and trimmed prior to construction of the phylogenetic tree, which was done with seaview BioNeighbor Joining Poisson software in the standard calibration mode. EvoMining (Cruz-Morales et al. 2016; Sélem-Mojica et al. 2019) and CORASON (Navarro-Muñoz et al. 2020) algorithms were used as previously. *S. roseus* ATCC31245 was obtained from the ATCC collection, and its genomic DNA was extracted using common protocols (Kieser et al. 2000) and sequenced at the genomic sequencing facilities of Langebio, Cinvestav-IPN (Irapuato, Mexico), using an Illumina MiSeq platform in paired-end format with read lengths of 250 bases and insert length of 800 bases. In total, 721 Mbp of sequence was obtained. The raw reads were filtered using Trimmomatic (Bolger et al. 2014) and assembled with velvet (Zerbino and Birney 2008), obtaining a 7.8 Mb assembly in 165 contigs with a coverage of 95 X and a GC content of 72 %. This assembly was annotated using RAST (Aziz et al. 2008), antiSMASH (Weber et al. 2015) and EvoMining (Sélem-Mojica et al. 2019). The genome includes 7,834,742 bp, and it was assembled in a total of 234 contigs and deposited in NCBI under the genome accession number NZ_LFML01000000.

### Construction of *Streptomyces* mutants and related *E. coli* strains

For the identification and characterization of livipeptin in *S. lividans* 66, a mutant deficient for the *Sli0883-5* genes was constructed. These genes were replaced by the apramycin resistance cassette *aac(3)IV* marker in-frame within a pESAC13-A construct. A 1.5 Kb region flanking the *Sli0883-5* genes replaced by the apramycin cassette was then amplified by PCR and cloned into the plasmid pWHM3, which contains a thiostrepton resistance gene. pWHM3 is an unstable *Streptomyces* vector that is lost after some rounds of cultivation of the transformed strain without selection (Vara et al. 1989). Double crossovers after integration of the pWHM3 *Sli0883-5:: aac(3)IV* construct were screened after apramycin resistance (50 μg/mL) and thiostrepton (25 μ/mL) sensitivity. The genotype of several transformants was confirmed by PCR, leading to *S. lividans* 66 ΔLvp used for experimentation. The synthetic livipeptin BGC, employing the *E. coli* usage codon, was obtained from GeneScript (New Jersey, USA). The design of the genetic construction includes restriction sites flanking each gene, as follows: *Nde*I-*Sli0883*-*Eco*RI-*Sli0884*-*Xba*I-*Sli0885*-*Bgl*II-*Hind*III. Different combinations of these genes were cloned into pAsk (Adams et al. 2014), resulting in six different plasmids (**Table 1** and **Supplementary Figure S5**).

The *Streptomyces roseus leupA* mutant was constructed following an insertional mutagenesis strategy using as parental strain ATCC 31245. For this, a *leupA* 640 bp fragment was PCR amplified and cloned into pCR2.1-TOPO using a TA cloning kit (Invitrogen, Carlsbad, USA). The resulting suicide plasmid, termed pLeupA, was introduced into *S. roseus* via protoplasts fusion (Kieser et al. 2000). The transformants were selected with kanamycin (50 μ/mL) and the genotype of the insertional mutant *S. roseus* LeuP1 was confirmed by PCR. The pESAC13-A leupeptin construct was obtained from a genomic library constructed at BioST, Montreal, Canada. The leupeptin BGC-containing cosmid pESAC_810, includes as insert the predicted leupeptin BGC. The genome of *S. lividans* 66 has been previously released (Cruz-Morales et al. 2013) and the genome of *S. roseus* is released here under the accession number NZ_LFML01000000.

### Culture conditions and sample preparation for metabolites analysis

Culture conditions for the induction of the livipeptin BGC were as previously (Cruz-Morales et al. 2013). Briefly, 50 mL shake flask cultures were inoculated with approximately 1×10^6^ fresh spores of *S. lividans* 66. Cultures were incubated at 30°C for 48h on R5 modified medium, without potassium phosphate, 0.2 g/L casamino acids and 200 mM MgCl_2_ (Kieser et al. 2000). The culture supernatants were collected for HPLC-MS analysis. For heterologous livipeptin production *E. coli* C41 Lvp strain was used to inoculate Luria broth (LB) cultures overnight and used as inoculum for 250 mL baffled flask with 50 mL of M9 medium (glucose 4 g/L added with L-arginine and L-phenylananine 0.5 g/L each) and incubated at 30°C and 200 rpm. The cultures were started at 0.1 OD_600_ and induced with 20 ng/µl of anhydrotetracycline. The culture was maintained for 24 h and the supernatant was collected for HPLC-MS analysis.

For production of leupeptin, *S. roseus* ATCC 31245 was grown on shake flask cultures containing a media designed by us, as follows: glucose 3 g, NH_4_NO_3_ 0.5 g, MgSO4 (7H_2_O) 0.5g, KCl 0.05 g, L-leucine 0.75 g, L-arginine 0.75 g, glycine 0.75 g, casamino acids 0.1 g, yeast extract 0.4 g per liter. Cultures were incubated at 30°C for 48 h prior to supernatant analysis. Heterologous leupeptin production was performed in shake flask cultures using *E. coli* DH10B bearing pESAC_810, in M9 minimal media (Glucose 4 g/L plus casamino acids 2 g/L) supplemented with apramycin (50 μg/ mL) for selection of the cosmid at 30°C for 48 h.

The cell extract of *E. coli* C41 pGroEL/GroES was prepared following a previously described method (Kigawa et al. 2004) with modifications. *E. coli* C41 pGroEL/GroES was inoculated in 50 mL de LB medium and grown at 37°C to an OD_600_ of 0.5 for induction with 20 ng/µl of anhydrotetracycline. After induction, the culture was grown at 20 °C for 12 h. Cells were harvested for (6000 rpm, 10 min, 4°C) and the pellet was washed three times by resuspension in S30 buffer (10 mM Tris-acetate buffer pH 8.2, 14 mM magnesium acetate, 60 mM potassium acetate, 1 mM dithiothreitol (DTT), 0.3 mM EDTA, 0.3 mM MgCl_2_) (Ortega et al. 2014) followed by centrifugation (6000 rpm, 10 min, 4°C). Cells were then resuspended in 1 mL of S30 buffer per gram of wet cells and lysed with a Sonicator. The cell lysate was centrifuged twice (13 000 rpm, 45 min, 4°C) and the supernatant was dialyzed four times against 50 volumes of S30 buffer (without DTT) using Amycon Ultra-15 (MerckMillipore) tubes for dialysis with a molecular mass cutoff of 10 kDa. The cell extract was then centrifuged (4 000 rpm, 10 min, 4°C) and the supernatant was frozen and stored in 1 mL samples at −80°C for future use.

### LC-MS analysis of purified metabolites

Supernatant of cultures and in vitro reaction mixtures were evaporated to dryness. The dry residues were dissolved in a 0.1 volume of HPLC grade H_2_O and injected into a C18 Discovery 504955 Supelco column with a particle size of 5 μm connected to a HPLC-Agilent 1200 equipped with a diode array detector and a fraction collector. The mobile phase comprised a binary system of eluent A, H_2_O, and eluent B, 100% MeOH. The run consisted of H_2_O/MeOH gradient (0-5 min: 0% B; 5-35 min: 10% B; 35-45 min: 100%. Differential peaks were detected between the wild-type and mutant strains (or empty vectors) by monitoring absorbance at a wavelength of 280 nm, and the selected fractions were collected for bioactivity assays and MS analysis in an ion trap LTQ Velos mass spectrometer (Thermo Scientific, Whatam, USA). MS/MS analysis of selected ions was performed with a collision energy of 20 eV.

### Protease inhibitory enzyme assays

Fractions collected in HPLC analysis were analyzed in vitro using a fluorometric assay with excitation at 340 nm and emission at 420 nm. The chromogenic compound Nα-Benzoyl-DL-arginine 4-nitroanilide hydrochloride (BApNA, Sigma-Aldrich) was used as substrate for trypsin and papain (Sigma-Aldrich T1005 and P3375, respectively), and leupeptin standard (Sigma, No. L2884) as positive control. HPLC peaks collected were dried in a vacufuge and then resuspended in milliQ water. The assay mixture for trypsin inhibition contained Tris-HCl (0.1 M pH 8) as reaction buffer, trypsin (0.05 mg/ml), substrate BApNA (0.1 mg/mL), leupeptin (0.001 mg/mL as positive control) or collected peaks (50 µL). Papain inhibition assay was done on a phosphate reaction buffer with DTT (100 mM), EDTA (60 mM), papain (0.01 mg/mL), antipain (0.01 mg/mL as positive control) or collected peaks (50 µL). The following reaction mixture in a final volume of 200 µL was used for the in vitro protease assays: E. *coli* cell free extracts or HPLC fraction (10 µl), HEPES pH 7.5 (100 mM) DTT (1 mM), L-lysine (10 mM), L-leucine (10 mM), L-Arginine (10 mM), L-Phenylalanine (10 mM), MgCl_2_ (10 mM), KCl (10 mM), ATP (5 mM) and NADPH (5 mM). The assay was incubated at 30°C for 5 h, centrifugated to remove insoluble material (13,000 rpm, 5 min, 25°C). In addition, the cell free extract (10 µL) was treated with RNase in the presence of CaCl_2_ (100 µM). The activity was calculated in percentage using as 100% the relative fluorescence units (RFU) of the proteases without inhibitors at the end of the reaction (20 min). The slope of the curve (initial rate) of the time progress of the reaction in each experiment was also calculated by triplicate.

## Supplementary Information

**Figure S1**. Complete phylogenetic tree of actinobacterial L/F transferases.

**Figure S2**. Complete EvoMining and CORASON analysis of leupeptin BGC.

**Figure S3**. High resolution MS spectral data of livipeptin.

**Figure S4**. HPLC-MS profiles of homologous and heterologous leupeptin.

**Figure S5**. Synthetic biology approach for the *cis* and *trans* expression analysis of livipeptin BGC.

## Acknowledgements

We thank the HPLC and MS Units of Cinvestav-IPN, Irapuato, for technical support. We also thank Pablo Cruz-Morales for strains construction and Nelly Selem-Mojica for bioinformatics support. This work was supported by Conacyt, Mexico (FINNOVA No. 214716), Laboratorios Liomont SA de CV and the UK Royal Society via a Newton Advanced Fellowship to FBG (NAF\R2\180631). CA was supported by a Conacyt postdoctoral fellowship (No. 730043).

